# CPlantBox, a whole plant modelling framework for the simulation of water and carbon related processes

**DOI:** 10.1101/810507

**Authors:** Xiao-Ran Zhou, Andrea Schnepf, Jan Vanderborght, Daniel Leitner, André Lacointe, Harry Vereecken, Guillaume Lobet

**Affiliations:** Agrosphäre (IBG-3), Forschungszentrum Jülich GmbH, Jülich, Germany; Simulationswerkstatt, Humboldtstraße 40, 4020 Linz, Austria; INRA, UMR 547 PIAF, F□63100 Clermont□Ferrand, France

## Abstract

The interaction between carbon and flows within the plant is at the center of most growth and developmental processes. Understanding how these fluxes influence each other, and how they respond to heterogeneous environmental conditions, is important to answer diverse questions in forest, agriculture and environmental sciences. However, due to the high complexity of the plant-environment system, specific tools are needed to perform such quantitative analyses.

Here we present CPlantBox, full plant modelling framework based on the root system model CRootBox. CPlantbox is capable of simulating the growth and development of a variety of plant architectures (root and shoot). In addition, the flexibility of CPlantBox enables its coupling with external modeling tools. Here, we connected it to an existing mechanistic model of water and carbon flows in the plant, PiafMunch.

The usefulness of the CPlantBox modelling framework is exemplified in four case studies. Firstly, we illustrate the range of plant structures that can be simulated using CPlantBox. In the second example, we simulated diurnal carbon and water flows, which corroborates published experimental data. In the third case study, we simulated impacts of heterogeneous environment on carbon and water flows. Finally, we showed that our modelling framework can be used to fit phloem pressure and flow speed to (published) experimental data.

The CPlantBox modelling framework is open-source, highly accessible and flexible. Its aim is to provide a quantitative framework for the understanding of plant-environment interaction.

## Introduction

Plants contribute for around 80% of the global biomass [1], and they strongly control land surface fluxes of water and carbon. Plant water uptake constitutes a major part of the evaporative flux at the land surface but its prediction is extremely variable and uncertain [2–4]. The same is true for the estimation of carbon fluxes [5,6]. As such, understanding the interplay between plant carbon and water flow and their environment is of importance to answer diverse questions in forest, agriculture and environmental sciences.

The flows of water and carbon in the plant are constrained by both local and overall structures [7–10]. Root architecture is known to have an impact on water uptake [11,12], while shoot structure has an impact on carbon assimilation [13–15]. From an entire plant perspective, root and shoot are tightly connected, forming a complex and dynamic continuum between water and carbon flow. For instance, water availability at the root level influences carbon unloading rate [16,17], while the stomata conductance directly affects root water uptake [18,19]. Knowing the connecting structure of both shoot and root is therefore needed to understand plants better.

At the organ scale, the different parts of the plant (root, stem, leaves, flowers and fruits) are connected by xylem and phloem vessels (fig. 1) [20,21]. Xylem vessels transport water [8,12,22,23], while phloem vessels translocate carbon [7,24–26]. The movement of water within the xylem vessels is typically explained by the tension-cohesion theory [27,28]. This theory states that transpiration at the leaf level creates a tension within the xylem vessels, which is transmitted to the soil-root interface and drives the water uptake from the soil. The carbon flow in the phloem continuum is explained by the Münch theory [29,30]. Briefly, Münch theory states that source organs (typically mature leaves and storage structures) load carbohydrates into the phloem sieve tubes. This strongly decreases the phloem water potential. As xylem and phloem vessels are tightly connected throughout the whole plant, this decreased osmotic potential in phloem will create a water flow from the xylem toward the phloem at source location (fig. 7**C** light green line). This in turn increases water pressure in the phloem vessels, leading to a flow toward sink organs (typically roots (fig. 7**C** light yellow line), young leaves, flower and fruits). [31,32]. Recent experiments have provided the first direct support to the Münch theory by direct measurement [33,34].

**Figure 1.**
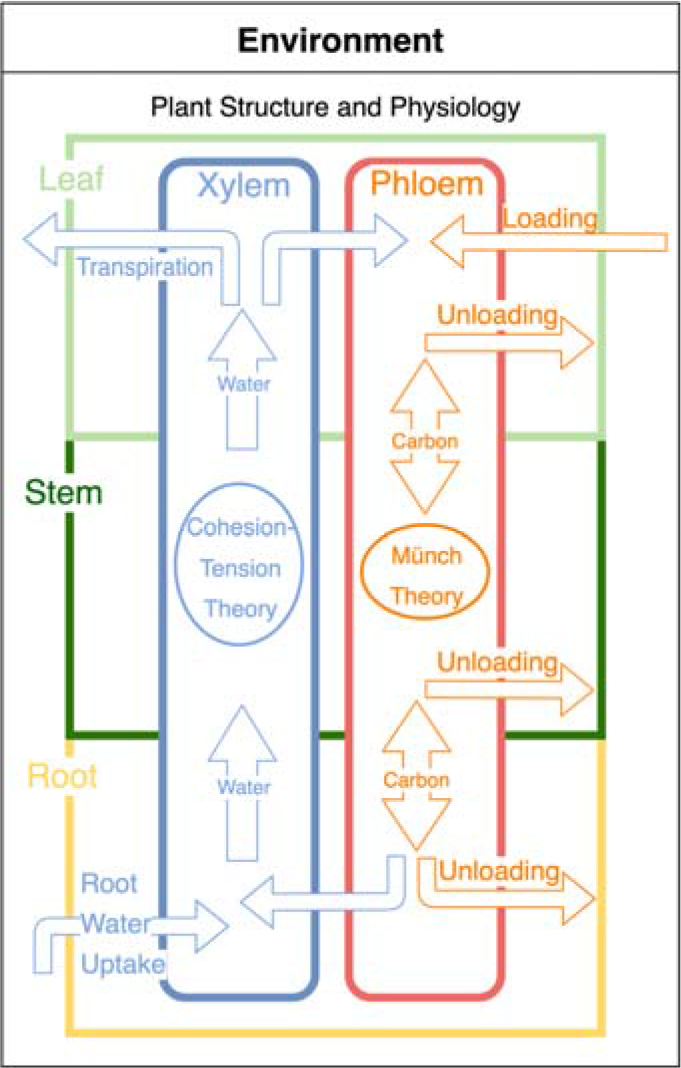
Structures and functions affect carbon and water flow in plants. Leaf and root contacts the environment. Xylem and phloem connect organs and exchange carbon and water.

In recent years, new phenotyping techniques [35–43] have enabled the precise measurement of plant structure with high temporal and spatial resolution. However, physiological parameters, such as pressure and flows in plants are still challenging to measure. For example, the first pressure measurement in phloem sieve tubes were conducted only recently with a success rate lower than 30% [33,44]. Another common issue with flow and pressure data is the fragmentation of the acquired data. In other words, as these techniques are complex to conduct, data can only be acquired on specific organs and at specific times, which makes it difficult to structurally understand the processes. More comprehensive and quantitative studies are therefore needed to better understand the complex dynamics between the water and carbon flow within the plant, in response to heterogeneous environments [45,46].

Recently, modelling tools have been proven very useful to study water and carbon flows in plants and to analyze environmental controls on these fluxes [16,47]. In particular, Functional-Structural Plant Models (FSPMs) have a long history of simulating water or carbon flows [48–56]. Table 1 lists the most recent FSPMs simulating either the full plant structure (both root and shoot), or water and carbon flows. Among these, only a handful of 3D full plant structure models (with both 3D topology and 3D geometry) exist [57–59]. Meanwhile, only two existing models were designed to simulate carbon and water flow simultaneously [60,61].

**Table 1.** Overview of recent 3D whole plant (shoot and root) FSPMs

| Model Name | Authors | Structure | Species | Flow(s) | Availability | Interface |
| --- | --- | --- | --- | --- | --- | --- |
| GRAAL<sup>1</sup> | [57] | 3D full plant | Maize | Water | - | GUI |
| PlaNet-Maize<sup>1</sup> | [59] | 3D full plant | Maize | Water | Open source | GUI, Web interface |
| Greenlab | [50,67,68] | 3D shoot and 0D root | Generic | Water or carbon | Upon request | GUI |
| .-<sup>2</sup> | [60] | 3D full plant topology | Small plants | Water and carbon | - | - |
| L-Peach | [69] | 3D full plant topology | Peach | Carbon | - | - |
| AmapSim | [70] | 3D shoot or root | Generic | Water or carbon (by coupling) | - | - |
| Piaf-Munch<sup>2</sup> | [61] | 3D full plant topology | Small plants | Water and carbon | Upon request | GUI, Command Line. |
| .-<sup>2</sup> | [58] | 3D full plant topology | Generic | Water | Upon request | - |
| CN-Wheat | [71,72] | 3D shoot, 0D root | Wheat | Nitrogen and carbon | - | - |
| This work<sup>1</sup> |  | 3D full plant | Generic | Water and carbon (by coupling) | Open source | GUI, iPython notebook, Web interface |

<sup>1</sup> Models that has both 3D topology and 3D geometry.<sup>2</sup> Model which can simulate interactions between carbon and water.

We distinguish three approaches to model carbon distribution within the plant. The first approach prescribes allocation rules of assimilates between the different plant organs. The total pool of carbon is divided between different organs, which adjust their growth accordingly [62,63]. Usually, models using such approach do not need a fast-computational method to distribute the carbon. A second approach uses detailed mechanistic relationships to simulate carbon (and sometimes water) flow within a simplified structure. These lumped models often only represent the plant as a small set of objects (fig. 2**B**) [25,64]. Finally, a third approach resolves carbon and water flow within a 3D structure based on mechanistic relations between the different organs [7,60,61,65]. Although these models can be computationally very intensive, they open the way to more complex representation of the plant-environment system.

**Figure 2.**
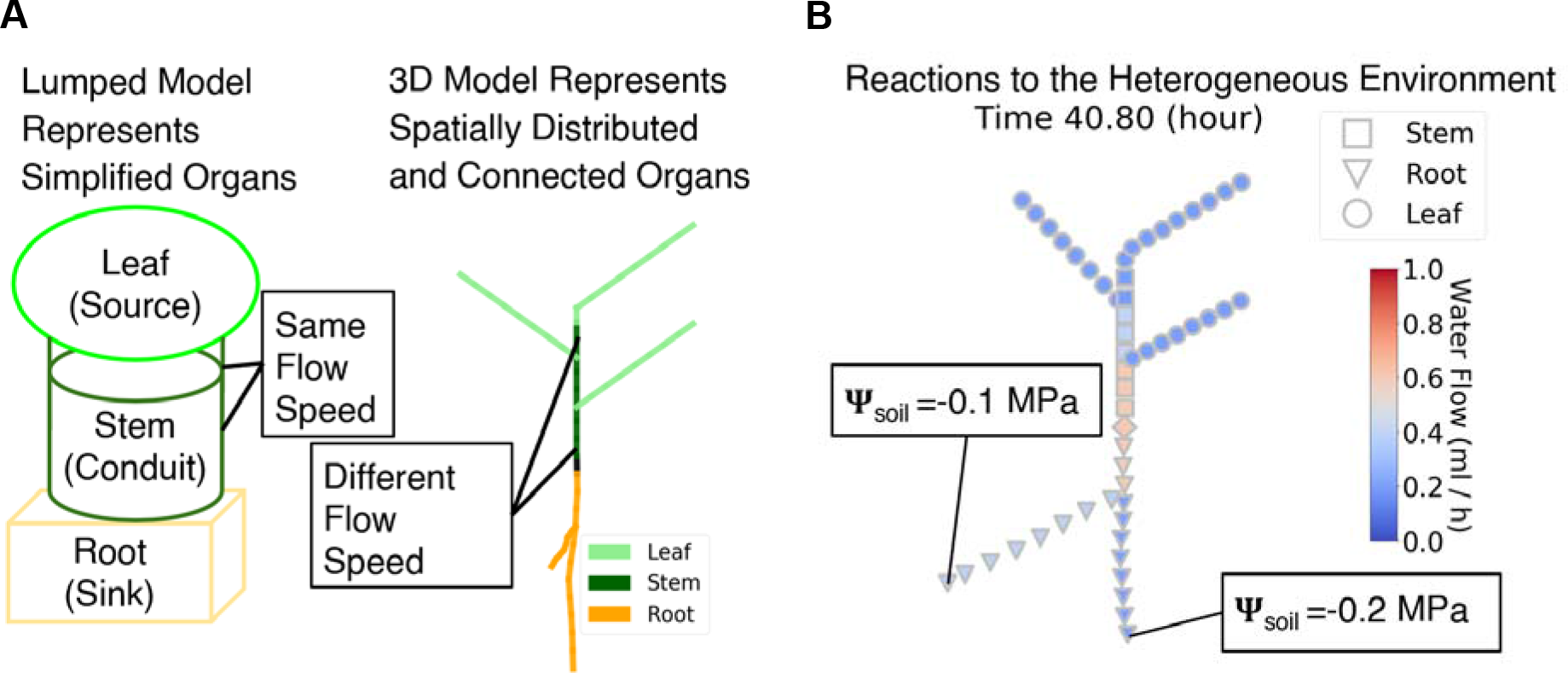
**A: T**he flow rate in lumped model depends only on the radius of the segment. In 3D models, branching structures affect flows as well. **B:** On 3D structures, the heterogeneous soil water potential affect xylem water flows. Root in wet soil has more water flow compared to root in dry soil.

Here we introduce a new framework for functional-structural whole plant modelling, which is based on the structural model CPlantBox. The novelty of CPlantBox is twofold. Firstly, CPlantBox can simulate the full plant structure at vegetative growth as a single connected network of organs (both root and shoot). Secondly, the model was coupled with a mechanistic model of carbon and water flow in the plant, PiafMunch [61,66]. The coupling enables fast simulations on large or complex plant structures, which was difficult to achieve before (PiafMunch uses manually defined plant architecture). Previously, PiafMunch is already able to simulate all 3D plant topology. Now, by coupling with CPlantBox, an additional 3D geometry layer is added to PiafMunch. Here we demonstrate the capabilities of the coupled model to generate a variety of plant structures and to reproduce realistic water and carbon flow behaviours.

## Material and methods

### Description of CPlantBox

CPlantBox is an extension of the model CRootBox [73]. CRootBox is a fast and flexible FSPM focusing on root architecture and root-soil-interaction. We took advantage of the object-oriented structure of CRootBox to add new modules to represent the different shoot organs (fig. 3**B**). The main extensions in CPlantBox are:

- CPlantBox can simulate realistic plant shoots and roots as a single connected network. The output can be coupled with water and carbon flow simulations (fig. 3**C**, **D**).
- As we move from root simulation to a full plant simulation (fig. 3**B**), more complex relationships between the different organs have been included in the model. For instance, roots can now grow from seed, roots or shoot organs (fig. 3**B**).
- The input parameter files are now XML-based (fig. 3**A**). Comparing to plain-text parameters, XML increases the robustness, flexibility (more parameters for the shoot) and readability. Backward compatibility with previous parameter file (from CRootBox) was insured.

**Figure 3:**
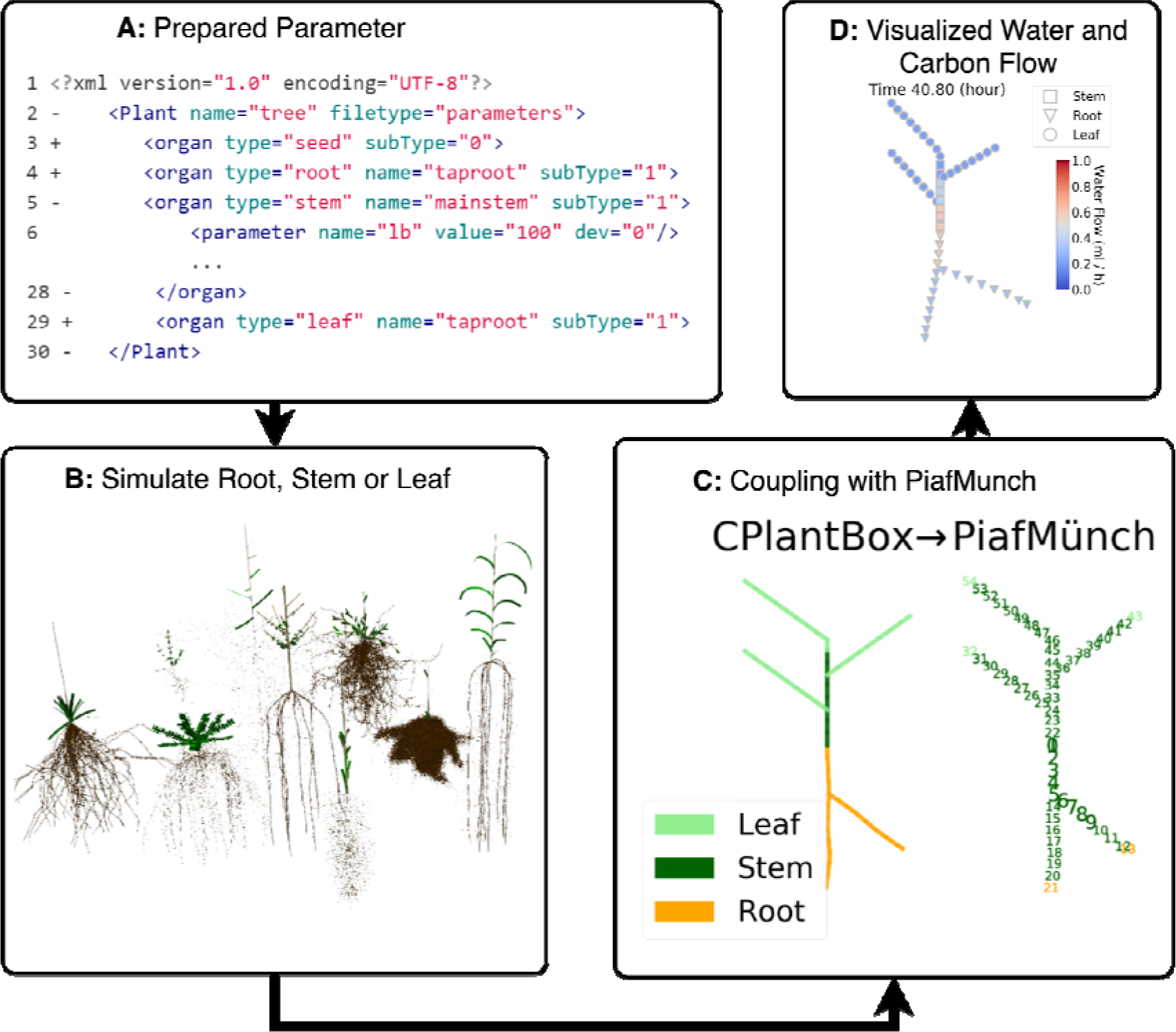
Flow chart of four major steps in CPlantBox-PiafMunch coupling. **A**: Input parameter example (organ parameters with “+” sign means it is collapsed). **B**: Output of CPlantBox is visualized. **C**: Coupling between CPlantBox and PiafMunch, where output of CPlantBox can be the input file of PiafMunch. **D**: Water flow output example from the CPlantBox-PiafMunch coupling.

The output of CPlantBox is a 1-dimensional network (with 3D geometry coordinates) (fig. 3**C**), each node of the network has 3D (3-dimensional) coordinates and other properties, such as an *organtype* (which indicates if it belongs to root, stem or leaf), a radius (or width) and a water potential. The output format of the structure includes RSML [74], VTP [75] and PiafMunch input format. After the plant structure is simulated (fig. 3**B**, potentially thousands of segments), an input file for the PiafMunch can be generated. Afterwards, the PiafMunch can be called by CPlantBox to read the input file, run simulation and generate the output files. At the end of the simulation, output files can be interpreted and visualized by the framework.

The current coupling is done by file exchange and command line automation. Simulating the carbon and water flow within a 300-segments plant for 100 hours growth time takes around 1 minute (dependent on parameter setting) on a laptop (CPU: Intel Core i5-6300U 2.4GHz, RAM 8GB 2400MHz). We also created looped functions to run simulations in batch processes. Installation, preprocessing, post processing and visualization are exemplified in the following Jupyter notebook: github.com/Plant-Root-Soil-Interactions-Modelling/CPlantBox/blob/master/tutorial/jupyter/CPlantBox_PiafMunch_Tutorial_(include_installation).ipynb.

### Description of PIAF-Munch

#### Water and carbon flows in PiafMunch

In PiafMunch [61,66], cohesion-tension theory is a precondition of the Münch theory. The cohesion-tension theory says that xylem-water-flow is driven by differences in pressure water potential in the xylem. The water potential in general can be defined as the sum of partial water potentials: the gravimetric water potential: □_z_ (MPa), the pressure water potential □_p_ (MPa) and the osmotic water potential: □_o_ (MPa). According to the cohesion-tension theory, the pressure in the water can be smaller than zero such that a tension instead of a pressure is applied to a water body. It must be noted though that mostly the water pressure is expressed as the difference between the water pressure and the atmospheric air pressure. Following this definition, a positive pressure corresponds with a pressure in the water that is larger than the atmospheric pressure and a negative pressure with pressure that is smaller than the atmospheric air pressure. Following this definition, a water tension would correspond with a water pressure that is smaller than −10^5^ Pa. If dissolved substances can flow freely within the xylem and phloem tissues, a gradient in concentrations or corresponding osmotic water potentials will not drive a flow within these tissues. Thus, the volume flow between two connected xylem’s n^th^ and (n+1)^th^ segments (*JW*_xyl,n,n+1_ mL h^−1^) can be written as:

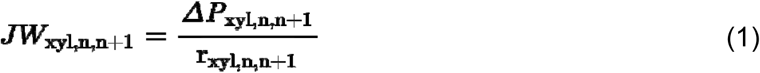

where r_xyl,n,n+1_ (MPa h mL^−1^) is the xylem resistance and *ΔP*_xyl,n,n+1_ = (□_z,xyl,n_+ □_p,xyl,n_)−(□_z,xyl,n+1_+ □_p,xyl,n+1_), *ΔP*_xyl,n,n+1_ is the difference between the sum of pressure and gravimetric potentials at the (n+1)^th^ and n^th^ segments respectively. Similarly, the volume flow between two connected phloem sieve tubes *JW*_st,n,n+1_ is:

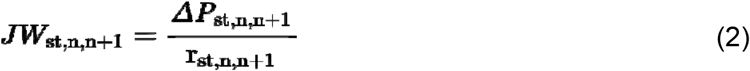

where r_st,n,n+1_ (MPa h mL^−1^) is the total sieve tube (include sieve plate) resistance between n^th^ and (n+1)^th^ segment and *ΔP*_st,n,n+1_ = (□_z,st,n_+ □_p,st,n_)−(□_z,st,n+1_+ □_p,st,n+1_), *ΔP*_st,n,n+1_ is the total potential difference between the neighbouring sieve tube segments, n^th^ and (n+1)^th^. At the n^th^ segment, the volume flow between the neighbouring xylem and phloem, which are separated by a semipermeable membrane, *JW*_lat,n_ can be written as:

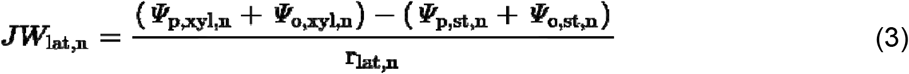

where □_p,xyl,n_ is the water pressure potential in xylem and □_p,st,n_ is the water pressure potential in sieve tubes, □_o,xyl,n_ is the osmotic water potential in xylem and □_o,st,n_ is the osmotic water potential in sieve tubes, r_lat,n_ is the resistance of the membrane between xylem and phloem. Here, we should notice that, at the source location, osmotic pressure drives the *JW*_lat,n_. But, at the sink location, the driving force is mainly the pressure water potential, because most osmotic water potential is removed by the unloading of carbon. The water mass balance of xylem is:

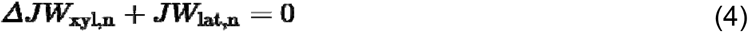

where Δ*JW*_xyl,n_ is the xylem water flux divergence over segment n, which can be written as Δ*JW*_xyl,n_ = *JW*_xyl,n,n+1_ − *JW*_xyl,n−1,n_. Similarly, the water mass balance of phloem can be written as:

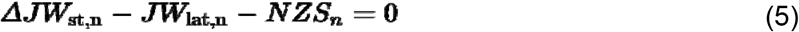

where Δ*JW*_st,n_ = *JW*_st,n,n+1_ − *JW*_st,n−1,n_ is the flux divergence over phloem sieve-tube. *NZS*_n_ is the non-zero sugar volume flow accompanying *JS*_lat,n_. At the source location, 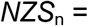 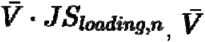 is the non-zero partial molar volume of sucrose, *JS*_loading,n_ (mmol h^−1^) is the loading rate from the source tissue (e.g. parenchyma) to the phloem at the n^th^ node. At the sink location, 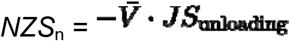, *JS*_unloading,n_ (mmol h^−1^) is the loading rate from the phloem to the sink tissue.

The mass balance of sucrose can be written as:

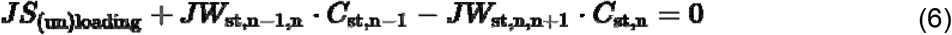

where, *JS*_(un)loading_ is the source (sink) term, *C*_st,n−1_ is the sucrose concentration in the (n−1)^th^ node, *C*_st,n_ is the sucrose concentration in the n^th^ node.

#### Comparison with experimental results

Recently, Knoblauch et al. experimentally tested the Münch theory on morning glory (*Ipomoea nil*) [33]. To validate the functions of CPlantBox-PiafMuch and estimated carbon loading/unloading speed, we decided to perform a re-analysis of their experimental dataset (measurements are shown in Table 2). In particular, we used one set of morning glory experimental data (left column of 7.5 m morning glory in Table 2) for calibration and another set (right column of 7.5 m morning glory in Table 2) for validation of our modelling system. We choose to use these datasets for different reasons. Firstly, the experimental measurements match almost directly both the input and output of the CPlantBox-PiafMunch model (fig. 4). Secondly, the authors performed a variety of experimental treatments, allowing us to parametrize our model on one experiment and validate on the other. Finally, the relatively simple architecture of the morning glory allowed us to focus our analysis on the resolution of carbon and water flow themselves.

**Table 2.**
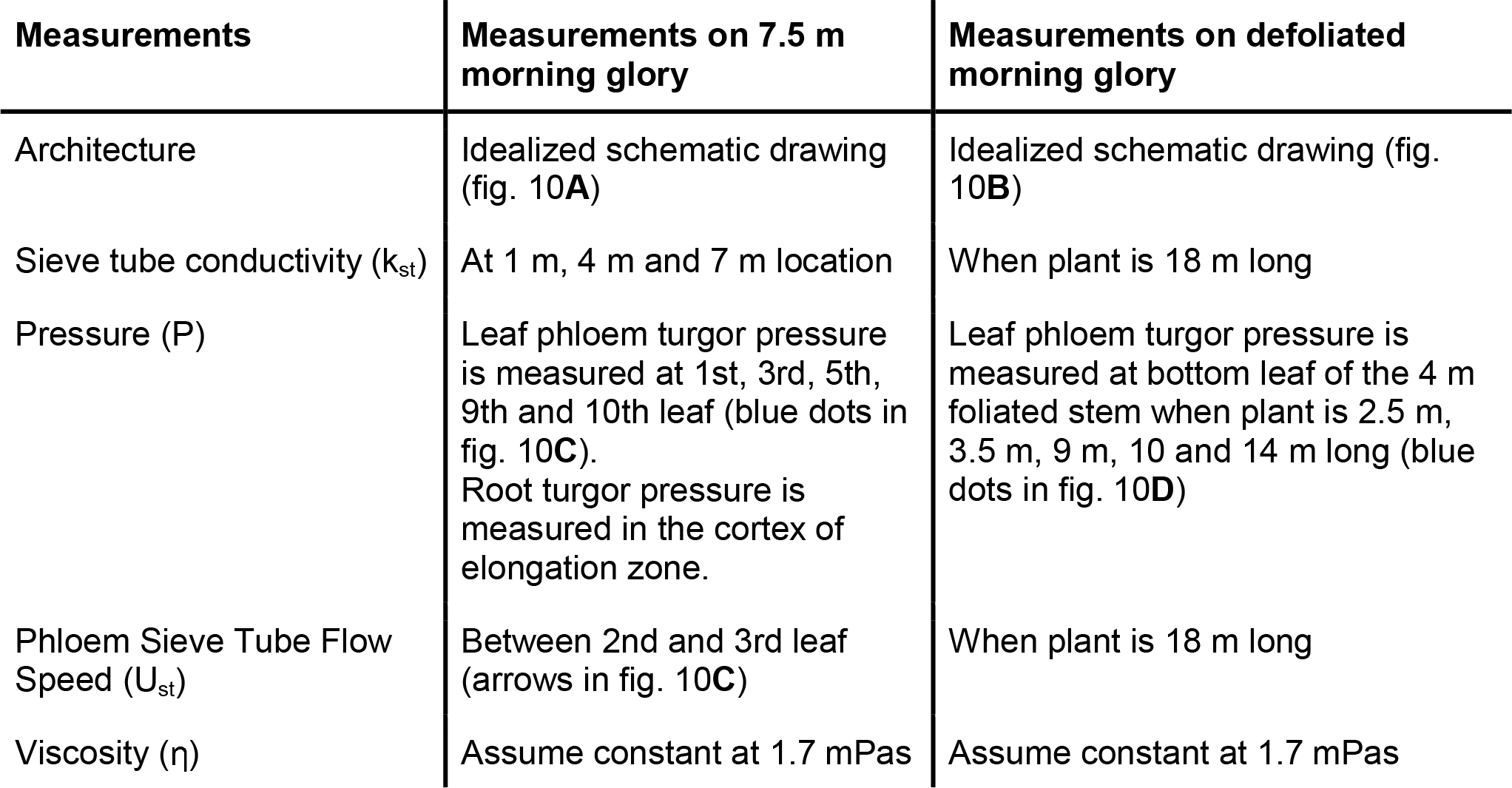
List of experimental measurements

| Measurements | Measurements on 7.5 m morning glory | Measurements on defoliated morning glory |
| --- | --- | --- |
| Architecture | Idealized schematic drawing (fig. 10A) | Idealized schematic drawing (fig. 10B) |
| Sieve tube conductivity ($k_{st}$) | At 1 m, 4 m and 7 m location | When plant is 18 m long |
| Pressure (P) | Leaf phloem turgor pressure is measured at 1st, 3rd, 5th, 9th and 10th leaf (blue dots in fig. 10C).<br>Root turgor pressure is measured in the cortex of elongation zone. | Leaf phloem turgor pressure is measured at bottom leaf of the 4 m foliated stem when plant is 2.5 m, 3.5 m, 9 m, 10 and 14 m long (blue dots in fig. 10D) |
| Phloem Sieve Tube Flow Speed ($U_{st}$) | Between 2nd and 3rd leaf (arrows in fig. 10C) | When plant is 18 m long |
| Viscosity ($\eta$) | Assume constant at 1.7 mPas | Assume constant at 1.7 mPas |

**Figure 4.**
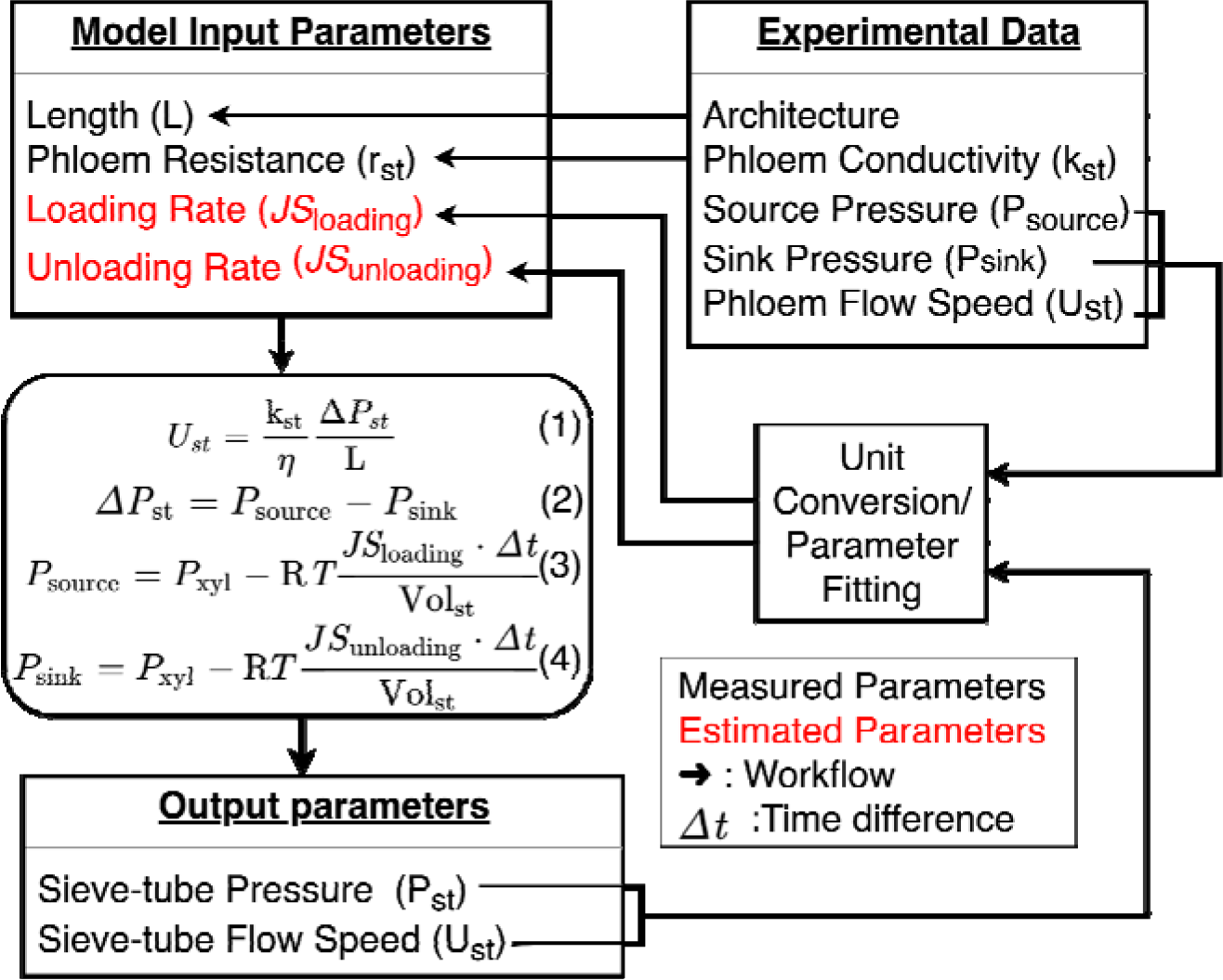
Measured parameters (in black text) and estimated parameter (in red text) in comparison with experiment. Length is calculated from simulated architecture (left side of fig. 10 **A, B**), which is based on the schematic drawing (right side of fig. 10**A**, **B**). r_st_ is converted from measured k_st_ (Supplement I, Equation 9). Pressure difference (Δ*P*_st_) is the phloem source pressure (*P*_source_) minus phloem sink pressure (*P*_sink_). R is the universal gas constant and T is the absolute temperature. *P*_xyl_ is the pressure of xylem at source or sink location, those pressures are calculated by carbon loading rate (*JS*_loading_) and carbon unloading rate (*JS*_unloading_). Arrows highlight our modelling workflow. Δ*T* is the time difference, Vol_st_ is the sieve tube volume. Units are in Table 3 and Supplemental material 1. The fitting of loading and unloading rate to source phloem pressure and flow rate are shown at the end of supplemental material 1.

**Table 3:** Literature values and simulation parameter values used in Example 2 and 4

| Parameter | Symbol | Type | Experimental Range | Unit | Small Plant | 7.5 m Plant | Defoliated Plant |
| --- | --- | --- | --- | --- | --- | --- | --- |
| Xylem Resistance | $r_{xyl}$ | Input | 0.011 to 3.3 by Stem [82] | MPa h mL<sup>-1</sup> | 2.08 by Stem<br>$6.9 \times 10^{-2}$ by Segment (0.25cm) | 0.081 by Stem 0.0005 by Segment (5cm) | 0.071 to 0.191 by Stem 0.0005 by Segment (5cm) |
| Phloem Resistance | $r_{phl}$ | Input | 40 to 90 [33] | MPa h mL<sup>-1</sup> | 70 by segment (0.25 cm) | Supplement Material I fig. 13 | Supplement Material I fig. 13 |
| Transpiration | - | Input | 0 to $1 \times 10^6$ m<sup>-2</sup> [83,84] | mmol h<sup>-1</sup> | 0.2 or diurnal | 0.2 | 0.2 |
| Soil Water Potential | - | Input | -0.2 to -0.8 [8,12] | MPa | -0.6 | -0.6 | -0.6 |
| Loading Rate | $JS_{loading}$ | Input | -- | mmol h<sup>-1</sup> | 0.0007 on each source segment | $6.4 \times 10^{-6}$ on each source segment | $6.4 \times 10^{-6}$ on each source segment |
| Unloading Rate | $JS_{unloading}$ | Input | -- | mmol h<sup>-1</sup> | $0.00028 \times C_{st}^1$ | $0.012 \times C_{st}^3$ | $0.012 \times C_{st}^3$ |
| Xylem Pressure | $P_{xyl}$ | Output | -0.2 to -8 [78] | MPa | 0 to -0.7 | 0 to -0.7 | 0 to -0.7 |
| Phloem Pressure | $P_{phl}$ | Output | 0 to 1.44 [85] | MPa | 0 to 1.8 | 0 to 1.4 | 0 to 1.8 |
| Xylem Water Flow Rate | $JW_{xyl}$ | Output | 0 to 3.6 [77] | mL h<sup>-1</sup> | 0.20 to 0.62 | 0 to 1.2 | 0 to 1.2 |
| Phloem Solut Flow Rate | $JW_{st}$ | Output | -0.02 to 0.02 [31,86] | mL h<sup>-1</sup> | -0.001 to 0.0006 | -0.002 to 0.0006 | -0.002 to 0.0006 |
| Phloem Carbon Flow Rate | $JS_{st}$ | Output | -- | mmol h<sup>-1</sup> | 0 to 0.00022 | 0 to 0.00022 | 0 to 0.00022 |

<sup>3</sup> $C_{st}$ : Carbon contraction in sieve tubes (details in Supplemental material 1, Equation 11).

In a first experiment (that we used for the parameterization of our simulations is shown in Table 2 and illustrated in fig. 10**A**), the authors measured the permeability of sieve tubes at three locations (1 m, 4 m and 7 m) on a 7.5 m tall morning glory (referred to as 7.5 m plant in following text). The phloem pressure and phloem flow rates were also measured in the same plant.

In a second experiment (that we used for validation shown in Table 2 and illustrated in figure 10**B**), another morning glory plant was continuously defoliated except for the top four meters (this plant is referred to as defoliated plant). When the defoliated stem was 2.5 m, 3.5 m, 9 m, 10 and 14 m long during its growth, the pressure of the bottom leaf was measured.

Details about the exact data transformations performed between the experimental measurements and the model parameters can be found in the Supplementary material 1.

## Results

In the following section, we presented four examples obtained with the CPlantBox-PiafMunch. Structurally, we present how CPlantBox can represent a wide variety of plant architectures. Functionally, we evaluated the quality of the water and carbon simulation of a three-leaf-two-root plant under either homogeneous or heterogeneous environments. In the end, we made a quantitative comparison between simulations and experimental data.

### Example 1: Simulation of contrasted plant architectures with CPlantBox

Plants can display a variety of forms and architectures, both above and below ground. [76]. Stem branching patterns are important factors determining the above-ground architecture of plants. fig. 5**A** shows an example of three branching patterns generated by CPlantBox using different parameter files. A second important determinant of the above-ground architecture is the arrangement of the leaves on the stems. fig. 5**B** includes three leaf arrangements also created by only changing single input parameter. By combining different branching patterns and leaf arrangements, we extended existing CRootBox outputs [73] into full plant architectures (fig. 6). It is worth mentioning here that each unique structure is obtained solely by changing the input parameter files. The source code itself is not modified.

**Figure 5:**
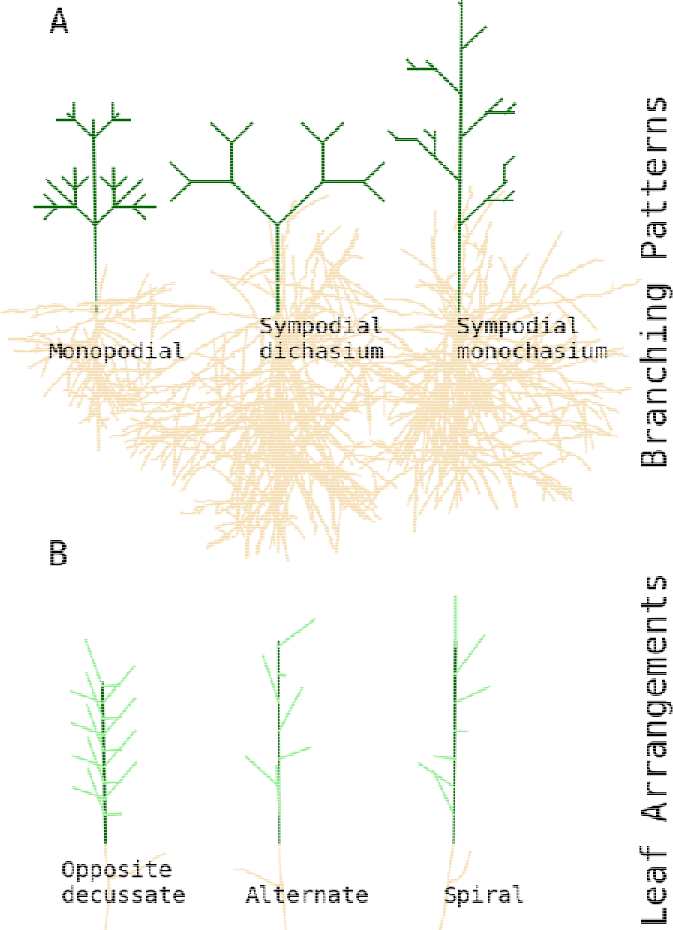
**A**: Simulated stem branching pattern simulated by CPlantBox; **B**: leaf arrangements simulated by CPlantBox

**Figure 6:**
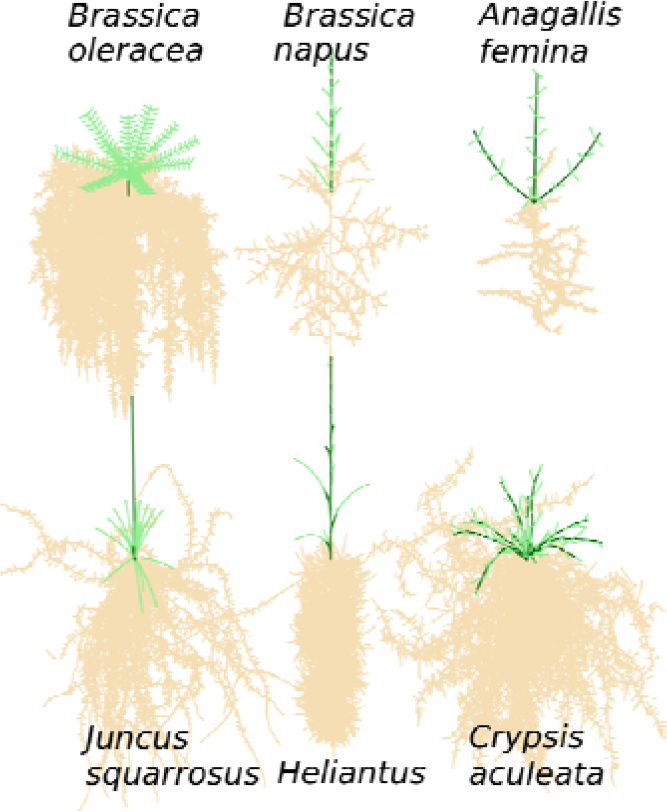
Simulated whole plant structures. *Brassica oleracea, Brassica napus, Anagallis femina, Juncus squarrosus, Heliantus* and *Crypsis aculeata*.

### Example 2: Simulation of water and carbon flow with the coupled model CPlantBox - PiafMunch

We created a smaller plant (it has three leaves and two roots) to simulate carbon and water flow (figs. 3**C** and 3**D**). The input and output parameters values were collected from various sources from the literature (summarized in Table 3)[77]. The simulated values of xylem pressure, flow rate and hydraulic conductivities are within the range of literature values. For example, xylem can sustain flow under pressure between −2.0 to −8.0 MPa, before losing 50% of its conductivity [78]. The simulated xylem water flow rate is typically around 1 ml h^−1^.

The transpiration rate on each leaf was set to mimic diurnal flow patterns. We set the transpiration rate to 0.2 mmol h^−1^ (0.0036 ml h^−1^) per leaf during daytime (from 5:00 to 17:30), and to 0 at nighttime (from 17:30 to 5:00 the next day). As shown in fig. 7**A** pressure is decreasing from root to leaf. Xylem flow during the day is caused by transpiration, and the phloem flow going back to xylem caused the xylem flow at night (which are lower than 0.0005 ml h^−1^, can be visible when zoom in fig. 7**B**). There are water moving from xylem to phloem at the source. According to Münch theory, the carbon loaded into the phloem will reduce water potential, so the water crosses the membrane, moves from xylem to phloem. Therefore, phloem carbon flow rates are affected by the diurnal xylem water flow (fig. 7**D**).

**Figure 7.**
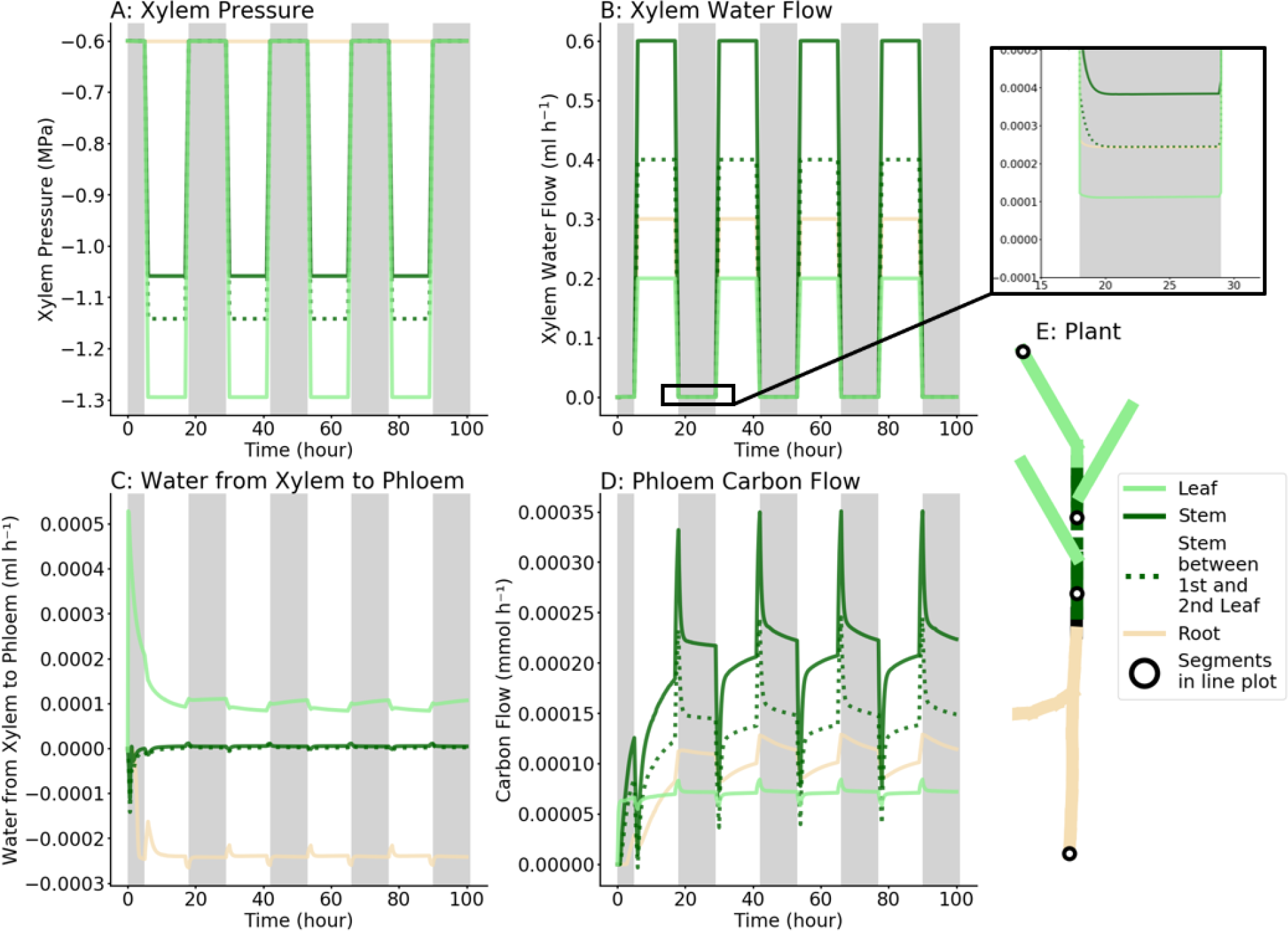
**A:** Transpiration creates xylem pressure gradients during the day, whereas the pressure remains comparatively stable during the night. **B**: The pressure gradient caused xylem water flow during daytime, the water flow at night is low, it is the water flow from phloem to xylem. **C**: The water flows from xylem to phloem at source location, whereas it flows from phloem to xylem at sink location. **D**: Phloem (sieve-tube) carbon flow fluctuations are caused by diurnal xylem water flow, the trend of changing is qualitatively consistent with previous studies [81]; **E**: Plant structure where the colors correspond to the flow figures from **A** to **D**, the dashed line shows the segments between the first and second bottom leaves. The flows going to circled segments are also shown as dashed lines in **B** and **D**. The pressure or flow exchanges of the circled segments are shown in **A** and **C.** The loading rate inside the phloem at source location is set to constant during both day and night, because starch is degraded to sucrose and then loaded into the phloem at night [79–81].

The loading rate into the phloem at source location is set to a constant value during both day and night. This is consistent with experimental data [79–81] as starch is degraded at night and the generated sucrose can be loaded into the phloem to sustain the flow.

### Example 3: Simulations of water and carbon flow in response to heterogeneous environments

Heterogeneous environments can have a large impact on plant growth and development. 4D FSPM can be used to simulate and visualize such environmental impact. To observe the effect of heterogeneous soil water availability on the carbon flow within the root system, we manually assigned two different soil water potentials at two root tips (bottom root in blue color with −0.2 MPa, upper root in red color with −0.4 MPa in fig. 8**A**). In fig. 8**B**, we observe a pressure difference between two roots, which causes hydraulic redistribution at night from the wet to the dry parts of the root system (fig. 8**C**). During the day, the water flow to the wet part is larger than the flow to the root in the dry soil. We also observed that the carbon concentration in the high water potential root is lower (red line in fig. 8**D**). In fig. 8**F**, we can see that total carbon flow in wet root is lower than the flow in the dry root.

**Figure 8.**
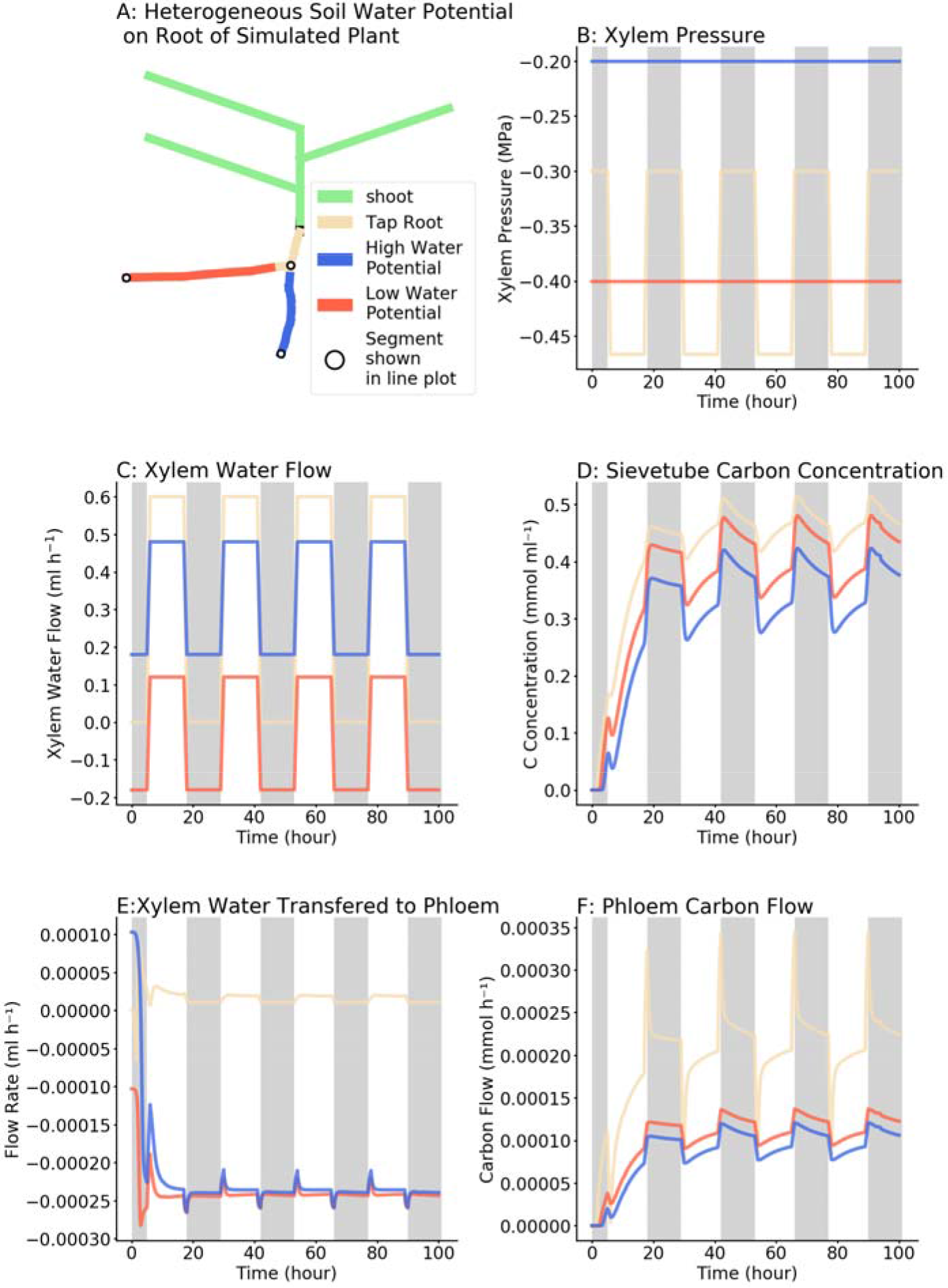
**A:** Soil water potential around the lower root (blue color) is higher than the upper root (red color). **B:** Pressure values at the boundary location are different according to the higher or lower soil water potential **C**: The bottom root (blue color, in wet soil) xylem, has higher water flow. The flow rate in upper root (red color) is negative at night. it means that, at night the water is coming out from the upper root to the soil, which is also called the hydraulic redistribution (in other words, the plant root system behaves as a pathway for water flow from wet to dry (or salty) soil areas). **D:** Carbon concentration in the dry (upper) root is higher than the wet root. **E**: Water flows from xylem to phloem at sink location only in the wet root (blue line) shortly at the beginning, then water flows from phloem to xylem in both roots at a similar rate. **F**: Carbon flow decreased in the high water potential (lower) root phloem.

Different temperature or developmental stages can also cause heterogeneous leaf transpiration rate. We assigned the 0.3 mmol h^−1^ (0.0054 ml h^−1^) transpiration on the top left leaf (higher transpiration leaf in fig. 9**A** with red color), 0.2 mmol h^−1^ (0.0036 ml h^−1^)transpiration on the right leaf (middle transpiration leaf in fig. 9**A** with green color), 0.1 mmol h^−1^ (0.0018 ml h^−1^) transpiration on the bottom left leaf (lower transpiration leaf in fig. 9**A** with blue color). In fig. 9**B** and **C** we can observe the pressure and flow gradient of three leaves at different transpiration rate. In fig. 9**D** and 9**E**, we can observe that carbon concentrations are different, but flows are slightly different between the different leaves as loading rate is kept constant. In fig. 9**F**, we can see that, when transpiration changes between day and night, the carbon flow in high transpiration leaf is more sensitive to the changes. However, the total carbon flow did not change significantly. In this example we keep the loading and unloading speed homogeneous and constant. It is because physiologically the starch degradation will compensate a temporal loss on the leaf level, just the same as the night carbon loading. Of course, the carbon loading will decrease in the long term, but it might not take effect in a few days.

**Figure 9.**
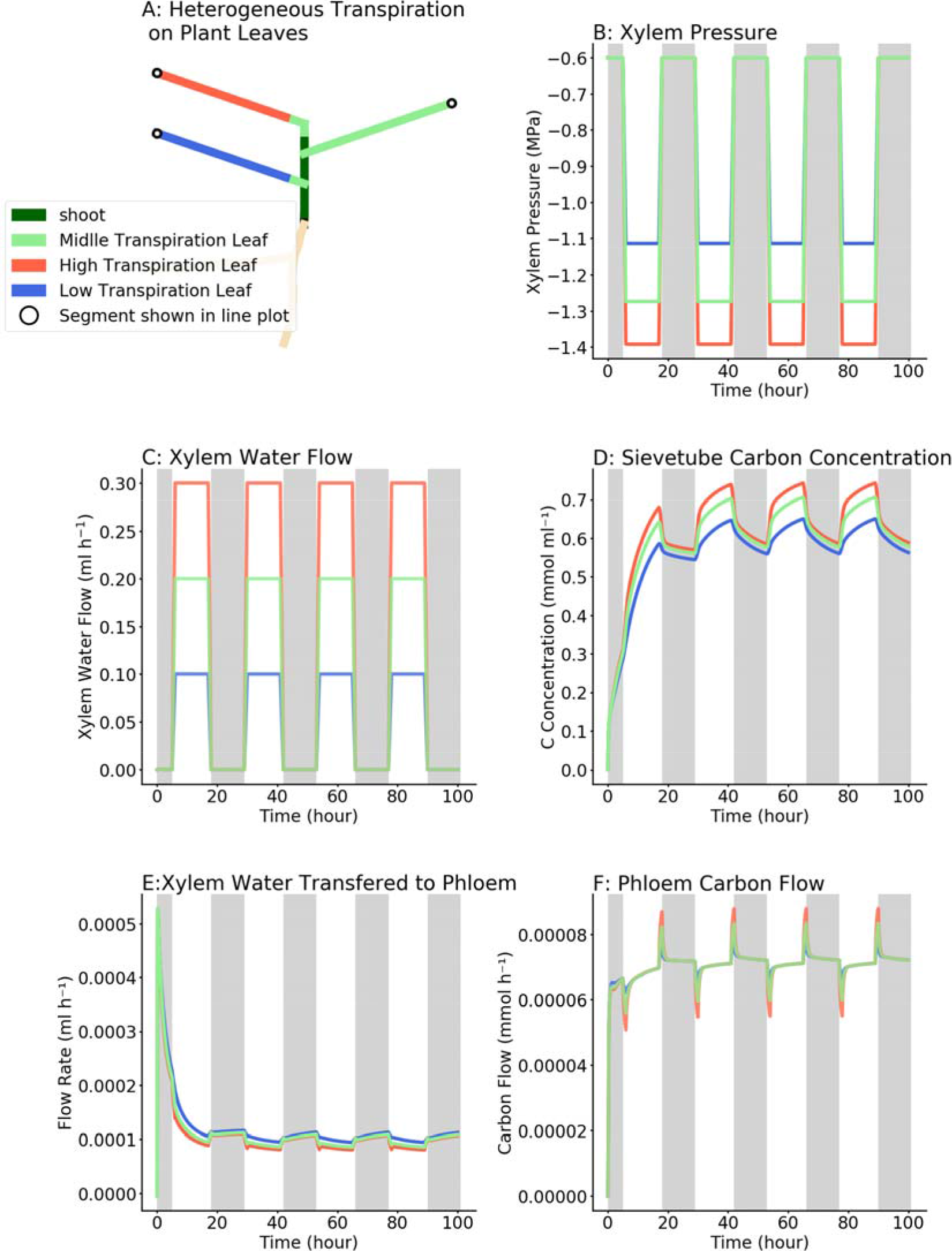
**A:** The lower left leaf (in blue) has lower transpiration rate and the top left leaf (in red) has higher transpiration rate. **B:** The pressure in each leaf is changed according to their transpiration rate. **C:** The water flow in each leaf is changed according to the transpiration rate. **D**: Carbon concentration on the higher transpiration leaf (red line) is higher. **B**: Water going from xylem to phloem at source location is lower at the higher transpiration leaf (red line). **F**: Carbon flow rates are the same at steady status, leave with higher water transpiration (red line) are more sensitive to the changes.

### Example 4: Comparison between experimental and simulated water and carbon flow in morning glory

To assess whether CPlantBox-PiafMunch was able to simulate realistic carbon and water flow values, we simulated experiments conducted with morning glory [33]. Three simulations were performed:

1. **Fitting parameters to measurements in literature (fig. 10 A, B, C**, **Table 2**): firstly we reproduced the plant structure based on schematic drawing. Then, we assigned the physiological parameters (phloem resistance) and used the estimated loading and unloading rate to fit the pressure and flow rate measured in phloem (fig. 4 and fig. 10**C** by method in Supplemental material 1, Table 4). In this case, we assumed that all leaves have homogeneous loading (which is also implicitly assumed in the experiment). **Testing fitted parameters (fig. 10 D**): we validated the loading and unloading rate on the 5 structures of the defoliated plant. The pressure matched the measurements.
2. **Conceptual experiment with individual loading/unloading on leaves (fig. 11 A, B, C, D**): The individual measured pressures are not consistent with the simulated pressure on each leaf, this could be a result of plant growth during the measurement or heterogeneous loading/unloading speed. Here we gave one possible example of the impact of heterogeneous loading/unloading speed on individual leaves.

**Figure 10:**
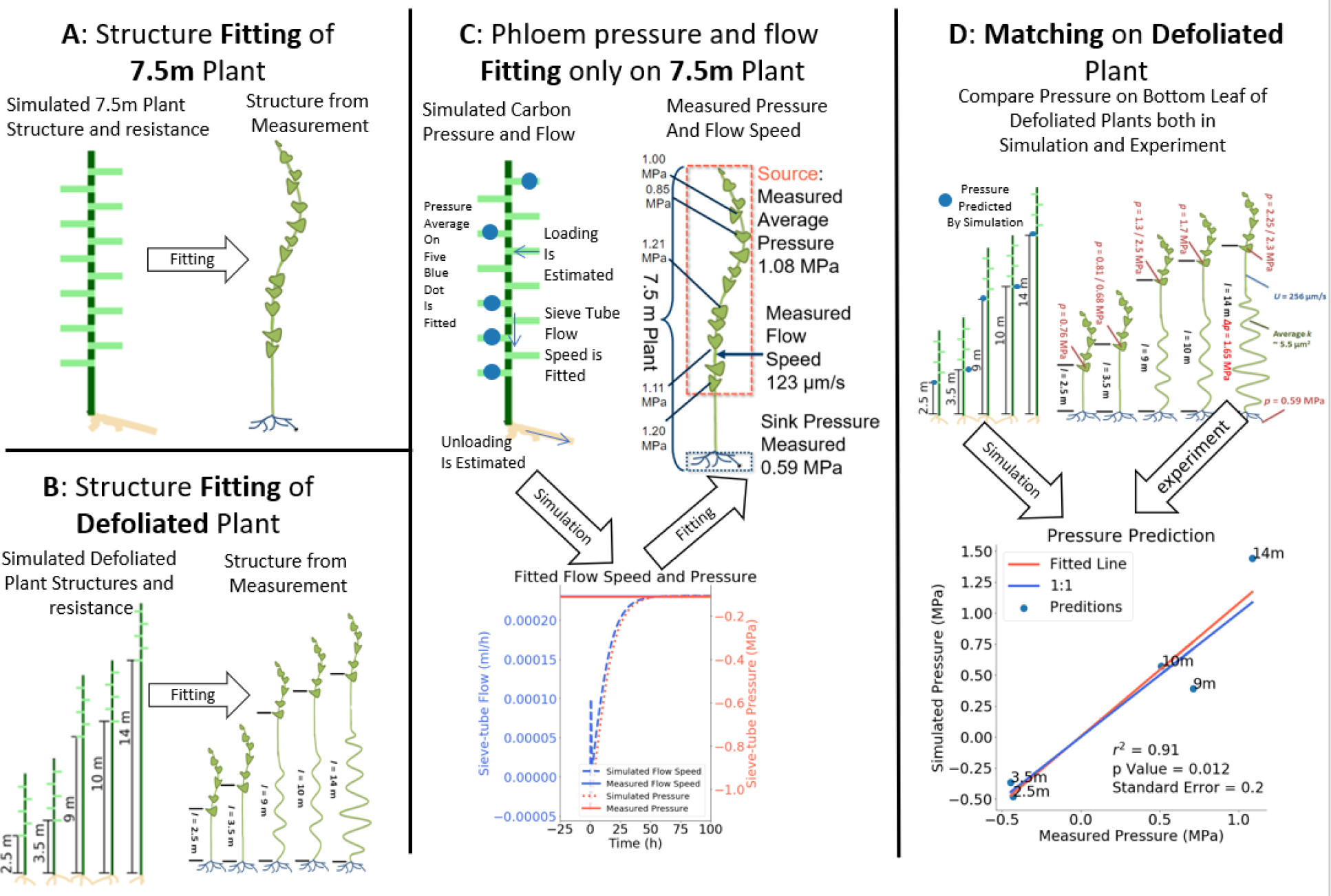
Steps of comparison between simulations and experiments. **A:** one *in silico* plant structure is created based on schematic drawing of the 7.5 m plant (resistance fitting can be found on Supplemental material 1, fig. 13). **B:** five *in silico* plant structures are created based on schematic drawing of defoliated plant. **C:** In phloem, physiological parameters such as source average pressure and sieve tube flow rate are fitted by applying homogeneous loading rate on each leaf, and homogeneous unloading rate at each root tip. Detailed parameters can be found in Table 2. **D:** By applying the fitted unloading rate and loading rate on 5 *in silico* plants, simulated pressure values match the measured values in experiment.

**Figure 11.**
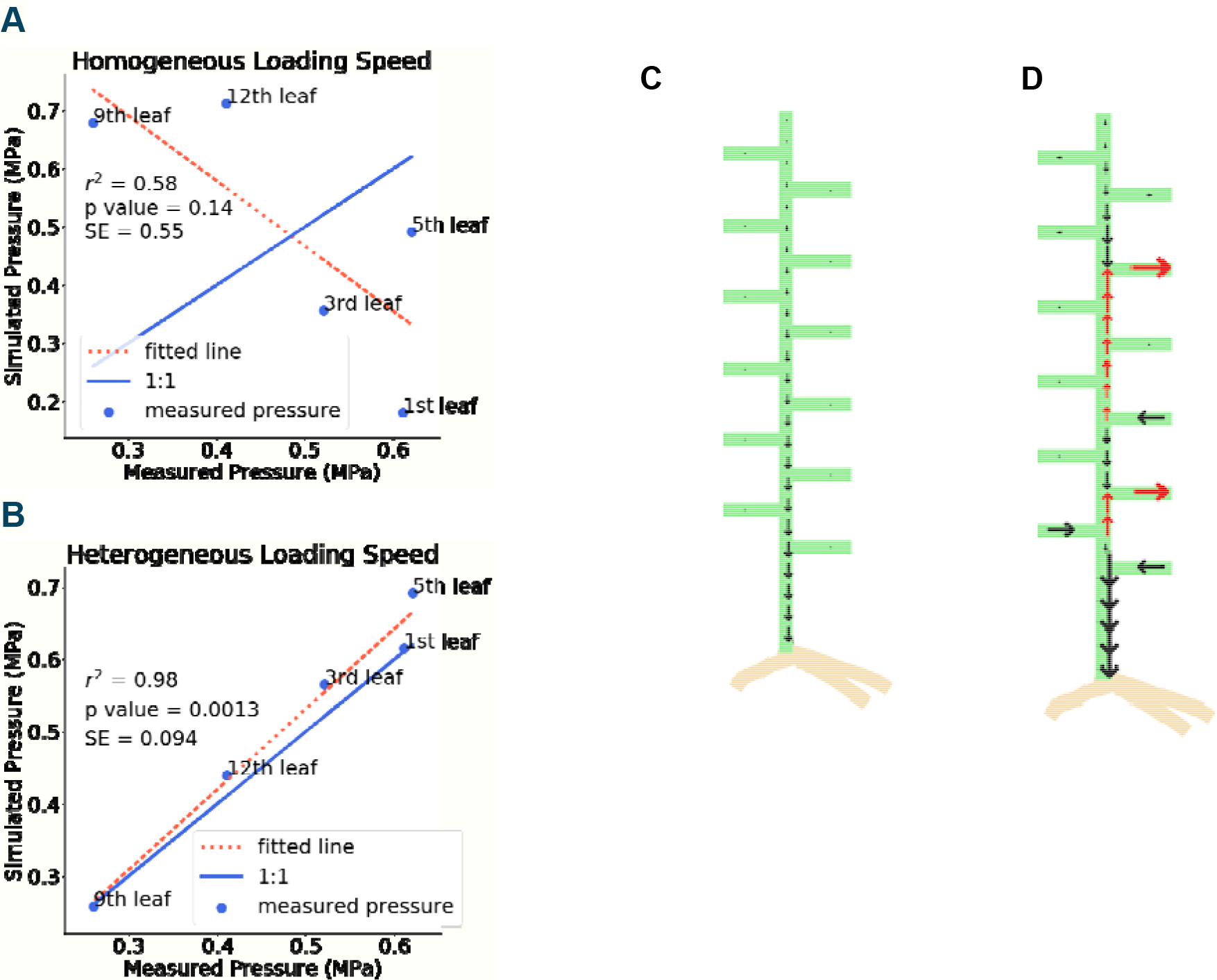
Comparison of simulations with (A) equal loading rate on all leaves to fit average pressure and (B) individual loading / unloading fitted pressure on each single leaf; **A:** Result for using homogeneous loading rate on each leaf; **B:** Fitted individual loading and unloading rate on each leaf; **C:** Flows are all heading to root when all leaves are sources with same loading rate (the scenario we used for parameter fitting); **D:** Flows directions changed (in red color) when the individual leave pressure are fitted (it is only one of the possible solutions, to show that we could change loading/unloading value on each source).

#### 4.1 Predicting carbon loading and unloading for contrasted morning glory shoot architectures

In order to simulate Knoblauch et al.’s experimental results on the morning glory, we created six virtual plants with contrasted architectures, based on the idealized schematics of the original paper. As described in Table 2, the reference plant was 7.5 meters long, with 12 homogeneously distributed leaves and one shoot tip(fig. 10**A**). The defoliated plants each had four leaves and one shoot tip near the apex of the stem, with different length for the defoliated section (fig.10**B**).

Regarding the physiological parameterization for the carbon and water flow simulation, in both the experiments and the modelling exercise, the morning glory was simplified to 1-source-1-sink system. Therefore, we assumed that the 12 leaves and the shoot tip are all homogeneous sources with the same carbon loading rate. The carbon unloading rates in the sinks were also considered homogeneous. Thus, we could create a 1-source-1-sink scenario as shown in fig. 10**C**, where the higher red dashed line box is the source and lower black dashed line box is the sink.

As shown in fig. 10**C**, we used the measured pressure and measured flow rate to find our initial input parameters, in particular the carbon loading and unloading rate. We estimated the corresponding loading and unloading rate using a looped least square fitting (lower part of fig. 10**C**, details are in Supplemental material 1, Table 4).

The carbon loading and unloading rate estimated on the 7.5-meter plant were then applied on the defoliated plants directly (Table 2 and fig. 10**D**). None of the parameters used in 7.5 m plant simulation were modified except the plant structure (fig. 10**B**). As shown in fig. 10**D**, we could see that the simulated pressure values in the sieve tubes were in good agreement with the experimental values.

#### 4.2 Studying source-sink relations at the organ level

In the previous section, the plant architecture was simplified to a 1-source-1-sink structure (fig. 10**C**), as in the experimental data analysis. In addition, we wondered if the model would be able to simulate the detailed relationship between different leaves inside the single source. It should be noticed that, as the large variance between the leaves might be caused by experimental variations, such detailed fitting might not be biologically relevant. However, it remains an interesting conceptual exercise, to test the flexibility and capabilities of our models.

First, we reused the calibration obtained on the 7.5 m plant (figs. 10 **A** and **C**). As shown in fig. 11**A**, it is obvious that the simulated pressure in each single leave does not fit the measured pressure. Therefore, we fitted each single leaf pressure by assigning independent carbon loading and unloading rate. We can observe that, when we reached a good fit on fig. 11**B**, the flow direction changed significantly compared to the flow when fitting the parameters globally (lower line plot of fig. 10**C**). Indeed, with the individual fitting, some leaves become carbon sinks instead of carbon sources. In fig. 11**D** the red arrows indicate a change on the carbon flow directions compared to the 1-source-1-sink scenario, as well changes in the total carbon loading (fig. 11**C**).

## Discussions

### CPlantBox generates full plant architectures

Historically, root and shoot models have been developed independently. Most models indeed focus on either part of the plant, representing the other one as a boundary condition. Some existing models are able to simulate both root and shoot, but for specific plant species [54,57,58,61,69]. Here, we presented CPlantBox, the first model, to our knowledge, able to represent both root and shoot, as a single network, for a variety of plant species (see example 1 in the Results section, and figs. 5 and 6).

CPlantBox was designed to be flexible and amenable for multiple plant studies. For the root part, CPlantBox inherited the flexibility of the model it was built upon, CRootBox. As CRootBox is able to generate any type of root architecture, so is CPlantBox. For the shoot part, we implemented several branching and leaf arrangement patterns. By combining these patterns, many types of shoot architectures can be simulated. Both root and shoot architectural parameters are defined into the model parameter file, making it easy to setup and reproduce.

### CPlantBox-PiafMunch simulates water and carbon flow in the full plant

We combined CPlantBox with a mechanistic phloem and xylem model, PiafMunch [61]. The coupled model allowed us to simulate water and carbon flow within complex full plant architectures, which was not possible before. In the results section, we demonstrated four examples. Example 1 is focusing on structures, while example 2 focuses on diurnal carbon/water flow compared to literature measurements. Example 3 shows that the combinations of structures and functions could reproduce literature experiments qualitatively. Example 4 firstly reproduce the experimental results then use conceptual experiments to prove that the heterogeneous environments (in Example 3) is helpful to explain experimental results.

In our simulation, we could observe a strong interplay between xylem and phloem flows. The diurnal transpiration patterns (the high peak in fluxes in the morning and the sharp decline when the light was stopped followed by an increase in flux during night) (fig. 7), consistent with previous experiment and modelling [81]. The low water potential in the xylem vessels during the day (as a result of the high transpiration rate) limits the water movement toward the phloem. During the night, as the stomata closes, the xylem water potential increases, leading to a higher water flow toward the phloem sieve tubes and a higher flow of carbon throughout the whole plant (fig. 7**D**). However, the higher night carbon flow might be caused by constant loading rate, whereas in some cases, the loading rate at night is reduced to 60% of the day value [87], so that the overall carbon flow may not be increased. In turn, the water flow from the xylem to the phloem induced a small upward water flow during the night, even in the absence of transpiration flux (fig. 7**B**). These results from our coupled models are comparable to previous published modelling results [61,66,88] and are consistent with experimental data (see Table 2 for details) [89].

### CPlantBox-PiafMunch considers the impact of heterogeneous environments

One of the main advantages of functional-structural plant models is their ability to explicitly consider the influence of heterogeneous environments (in space and time). In our third example, we used our coupled CPlantBox-PiafMunch modelling framework to simulate the influence of heterogeneous soil and atmospheric conditions on the carbon and water flows in the plant.

First, we imposed different soil water potentials to the different roots of our plant (fig. 8**A**). In response to this heterogeneity, we could make two main observations. Firstly, root water potential and water flow (fig. 8**B**, 8**C**) was directly influenced by the soil water potential. As the soil water potential decreases, the water flow in the xylem decreases. This is a well-known effect, observed both *in vivo* [90,91] and *in silico* [23,92]. Secondly, we observed that the carbon flow (fig. 8**F**) in the phloem was inversely correlated with the soil water potential. Indeed, our simulation results show that carbon flow is slightly higher in the portion of the root system in contact with a dry soil (red line in fig. 8**F**). This is due to the lower carbon concentration of root phloem in wet soil (blue line in fig. 8**D**). The lower carbon concentration in wet root phloem (blue line in fig. 8**D**) is a result of dilution by water from wet root xylem to wet root phloem along the root until the root tip (like tap root in light yellow color). At the root tip, the unloading rate is proportional to the carbon concentration. Thus, the flow rates of two split root are similar. Because the flow rates are similar, but concentration is lower in the wet root, the total carbon flow is lower in the wet root (fig. 8**F**). Again, this dynamic was observed experimentally for several plant species in split root experiment [93–95].

To simulate heterogeneous atmospheric environment, we imposed different transpiration rates to the different leaves of our plant (fig. 9**A**). Like water potential change at the sink location, the transpiration rate at the source location directly induced changes of xylem pressure (fig. 9**B**) and xylem water flow rate (fig. 9**C**). Heterogeneous transpiration rates on leaves are also observed *in vivo* and simulated *in silico* [96,97]. In fig. 9**D**, we could observe that the carbon concentration in phloem is increased, because in fig. 9**E** we could observe that in the high transpiration leaf (red line), less water is moving from xylem to phloem. This is because the water potential increases (red line in fig. 9**B**) and pressure drops (red line in fig 9**A**) in the high water potential leave. Thus, the final phloem carbon flow rate did not change at steady state (lines are aggregating in fig. 9**F**)

### Limitations of the model and future perspectives

In this paper we highlighted some of the capabilities of our new coupled model CPlantBox-PiafMunch. We have shown that we can simulate realistic water and carbon flow within a full plant structure. However, it is important to stress the current limitations and future developments of our model.

Firstly, all the simulations were done with static plants. At this stage, we did not explore the impact of the carbon distribution on the growth and development of the plant. In the future, we will explicitly connect the growth function in CPlantBox to the local carbon availability as prescribed by PiafMunch.

Secondly, in the presented simulations, the environment was static as well. In order to explore only the resolution of carbon and water within the plant, we did not connect our models to dynamic representations of the environment. Again, this will be done in the future, as we plan to integrate CPlantBox into the modeling framework DuMuX [98]. By doing so, we will be able to explore the feedbacks between the plant, the soil and the environment.

Finally, in the version of the models, carbon production is prescribed at the article level. Again, this what not an issue so far, as we wanted to explore the flow distribution only. However, in the near future, we plan to include leaf-level photosynthesis module [99,100], to be able to better represent the dynamic response of the plant to its changing environment.

## Conclusions

Models can be used as analysis tools for complex experimental setups. Experimental measurements of carbon and water flow can be challenging. Most available measuring methods are time consuming and destructive [33,44], preventing the continuous observation of these flow as the plant develops.

Here we have used our coupled models to reverse estimate hidden experimental parameters. For instance, by using measured carbon flow, phloem resistance and phloem pressure, we were able to give consistent estimates of carbon loading and unloading rate in the phloem, in the different plant organs.

More generally, this is a good example of using models as complex analysis tools. As experimental setup and biological questions become more and more complex, it becomes harder to interpret the results. Models such as CPlantBox-PiafMunch can help integrate such results and replace them into a whole plant perspective. Carefully using the model can then give access to additional parameters that were not available experimentally.

Exploring the interplay between the environment, the plant architecture, and the plant water and carbon flow is experimentally challenging. Measurements take time and are often destructive. However, functional structural plant models have been shown to be able to efficiently represent plant-environment interplay *in silico*.

Here we have presented a new whole plant framework, CPlantBox. We have shown that CPlantBox is able to represent a variety of plant architectures, both root and shoot. We also connected CPlantBox to a mechanistic model carbon and water flow, PiafMunch. The coupled model was able to reproduce realistic flow behavior in complex plant structures. We were also able to use the models to reproduce experimental data and estimate hidden experimental variables.

In the future, the model will be extended to more realistic growth behaviours, environmental changes and photosynthesis dynamics.

## Model and data availability

- CPlantBox is open source under GPL 3.0 license, available at https://github.com/Plant-Root-Soil-Interactions-Modelling/CPlantBox/tree/master/ and http://doi.org/10.5281/zenodo.3518068
- Model parameter files are available at: http://doi.org/10.6084/m9.figshare.9785396
- PiafMunch output of simulation example 2, 3, 4 can be found here: https://figshare.com/articles/Output_of_CPlantBox-PiafMunch_coupling/9971225

## Acknowledgements

We would also like to show our gratitude to Xavier Draye, Mathieu Javaus, Michael Knoblauch, Clément Saint Cast for sharing their pearls of wisdom with us during this research.

## Authors contributions

Writing – original draft: XRZ, GL; Writing – review & editing: JV, AL, HV, GL, XRZ; Conceptualization: GL, AS, HV, JV, XRZ; Software: XRZ, DL, AL, AS, GL; Visualization: XRZ, GL, JV

