## Supplemental File 1 for "CPlantBox, a whole plant modelling framework for the simulation of water and carbon related processes"

### Supplemental material 1

#### Variable and unit conversions

##### 1. Parameterization of architecture

The simulated plant structures are based on schematic drawings. The defoliated stem is clearly indicated from the schematic drawing. However, the number of leaves might not be exactly correct. Here, we can only assume that the schematic drawing could ideally represent the correct leaf numbers ratio across different structures.

##### 2. Parametrization of physiological parameters

The five variables in Hagen-Poiseuille equation (Equation (1) in fig. 4) were measured by Knoblauch et. al.. The length (*L*) is calculated based on architecture, viscosity (η) is assumed constant, resistance (r) is measured at three locations on the architecture. According to the Münch theory, when the other parameters stay unchanged, the flow speed (*U*) and pressure (*P*) value varies depending on loading and unloading rate (Equation (2), (3), (4) in fig. 4). Thus we could test different loading and unloading rate to fit the pressure and flow rate. Following variables and units are converted to a uniform format, which could be used in the models. The total sieve tube conductivity k_st_ (include bother sieve-tube and sieve plate) is measured by physical attribute of sieve-tube and sieve plate.

##### 3. From conductivity k_st_ (μm^2^) to resistance r (MPa h mL^-1^)

According to the anatomical structure of sieve tubes measured in the 7.5 m plant, the k can be calculated by Equation 9

| 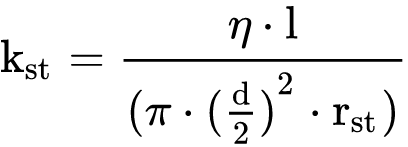 | (7) |
| --- | --- |

l is the metamer segment length, which is 0.05 meter long; η is the viscosity, which is assumed to be stable at 1.7 mPas; d is the diameter (radius×2) of sieve tube ; k_st_ (μm^2^) is the total sieve-tube conductivity (permeability). The resistance r_st_ (total resistance of the sieve-tube and sieve plate in one segment of phloem) at 1, 4, 7 meters are shown in fig. 13, where two linear equations are used to estimate all the resistance between each segment.

| 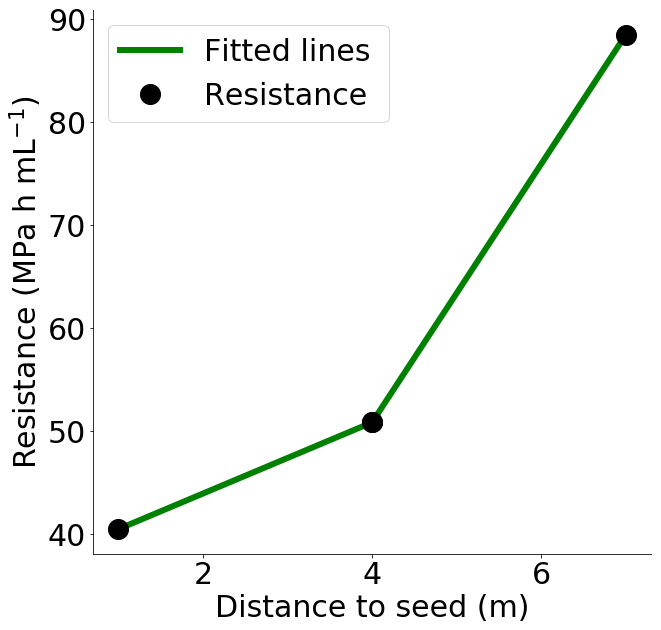 | **Figure 13:** Three black points are total sieve tube resistance (r_st_) calculated by Poiseuille’s law based on measured physical attributes of phloem ( diameter, length, sieve plate pore size etc. ). Green lines are the estimation of each segment resistance based on their distance to the seed. |
| --- | --- |

##### 4. Convert flow speed units µm s^-1^ to flow rate unit ml h^-1^

| 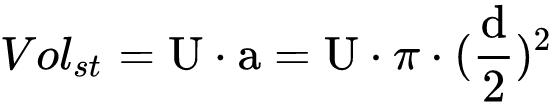 | (8) |
| --- | --- |

In Equation 8, V_st_ is the volumetric flow rate, U is the average flow velocity and a is the cross sectional area, d is the diameter (radius×2) of sieve tubes.

##### 5.  Estimating loading and unloading rate of 7.5 m plant

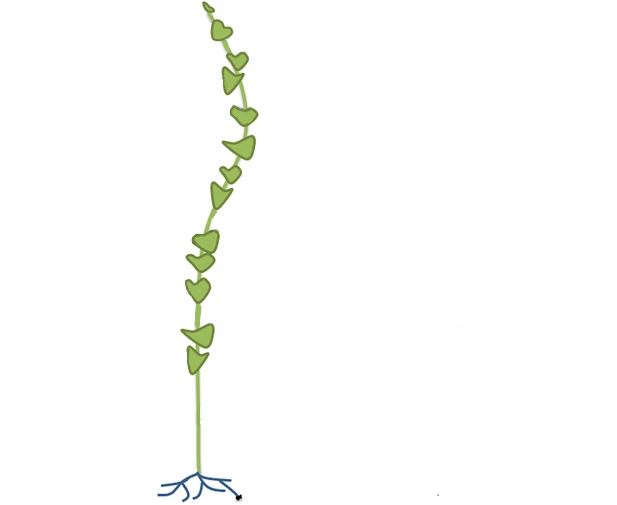

**Figure 14:** Schematic drawing of the 7.5 m plant.

According to the schematic drawing shown in fig. 14, we assume that all 12 leaves and one shoot tip ( in total 13 sources) of the 7.5 meter plant in fig. 10**A** are homogeneous distributed along the stem. Each leaf tip could load carbon at the same rate. We also set all sieve tube volume to 1.73 × 10^-7^ ml, which is the volume measured at 4 m position of the 7.5 meter plant. Loading and unloading rate are set by two Michaelis-Menten equations in Equation 9 and Equation 10.

| 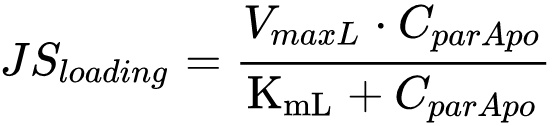 | (9) |
| --- | --- |

| 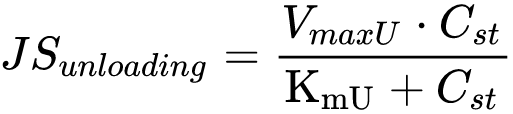 | (10) |
| --- | --- |

Where JS_loading_ (mmol h^-1^) is the loading rate, *V*_max_ (mmol h^-1^)_L_ is the maximal loading rate, K_ml_ is the Michaelis-Menten constant which should be the concentration when the reaction speed is half of the *V*_maxL_, *C*_parApo_ is the sugar concentration at parenchyma apoplas. *JS*_Unloading_ is the unloading rate, *V*_maxU_ is the maximal unloading rate, K_mU_ is the Michaelis-Menten constant which should be the concentration when the reaction speed is half of the *V*_maxU_ . *C*_st_ is the sugar concentration at sieve tubes.

In our simulation, the Michaelis-Menten equations are modified. For example, in the Equation 9 the K_mL_ is set to 1 × 10^-99^ (an extremely small number), so the *JS*_loading_ is equal to *V*_maxL_. In Equation 10, the K_mU_ is set to 1 × 10^99^ (an extremely large number), so *JS*_unloading_ is equal to *V*_maxU_ × *C*_st_, which means that the loading rate is a constant, while unloading rate change with the carbon concentration. After hand fitting, we could find that when *JS*_loading_ = 7.2716 × 10^-6^ mmol h^-1^, *JS*_unloading_= 0.0001672 × *C*_st_ mmol h^-1^.

#### Fitting loading and unloading rate to source phloem pressure and flow rate

According to Münch theory, a proportion of phloem pressure at source location is the osmotic water potential caused by the loaded sugar in phloem. At the same time, the increase of unloading rate could also cause the increase of phloem sap flow. Their relationship could be summarized in Table 4. Based on this relationship, we could build a loop function to estimate the loading and unloading rate.

**Table 4**. Relation between loading rate, unloading rate, source phloem pressure and phloem flow rate.

| Loading Rate | Unloading Rate | Source Phloem  Pressure | Phloem flow rate |
| --- | --- | --- | --- |
| **⭡** | **-** | **⭡** | **⭡** |
| **⭣** | **-** | **⭣** | **⭣** |
| **-** | **⭡** | **⭣** | **⭡** |
| **-** | **⭣** | **⭡** | **⭣** |
